# AKT1-FOXO4 AXIS RECIPROACLLY REGULATES HEMOCHORIAL PLACENTATION

**DOI:** 10.1101/2022.06.15.496110

**Authors:** Keisuke Kozai, Ayelen Moreno-Irusta, Khursheed Iqbal, Mae-Lan Winchester, Regan L. Scott, Mikaela E. Simon, Masanaga Muto, Marc R. Parrish, Michael J. Soares

## Abstract

Hemochorial placentation involves the differentiation of specialized cells called invasive trophoblast cells possessing the capacity to exit the placenta and invade into the uterus where they restructure the vasculature. Invasive trophoblast cells arise from a well-defined compartment within the placenta, referred to as the junctional zone in the rat and the extravillous trophoblast cell column in the human. In this study, we investigated roles for AKT1, a serine/threonine kinase, in placental development using a genome-edited/loss-of-function rat model. Disruption of AKT1 resulted in placental, fetal, and postnatal growth restriction. Forkhead box O4 (***Foxo4***), which encodes a transcription factor and known AKT substrate, was abundantly expressed in the junctional zone and invasive trophoblast cells of the rat placentation site. *Foxo4* gene disruption using genome-editing resulted in placentomegaly, including an enlarged junctional zone. AKT1 and FOXO4 regulate the expression of many of the same transcripts expressed by trophoblast cells; however, in opposite directions. In summary, we have identified AKT1 and FOXO4 as part of a regulatory network that reciprocally controls critical indices of hemochorial placenta development.

**SUMMARY STATEMENT:** Genome-edited rat models were utilized to investigate roles for AKT1 and FOXO4 in hemochorial placentation. AKT1 and FOXO4 possess reciprocal actions in regulating development of the hemochorial placenta.

## INTRODUCTION

The placenta is an extraembryonic structure essential for normal fetal development (**Maltepe & Fisher, 2015; Burton *et al*., 2016**). Placentas possess two main functions: i) transformation of the maternal environment to support viviparity and ii) regulation of the transfer of nutrients to the fetus (**Gardner & Beddington, 1988; Soares *et al*., 2018; Knöfler *et al*., 2019**). These specialized functions are ascribed to trophoblast cells, which differentiate along a multi-lineage pathway and are situated within specific compartments of the placenta (**Gardner & Beddington, 1988; Soares *et al*., 2018; Knöfler *et al*., 2019; Aplin & Jones, 2021**). Placentas come in different shapes, sizes, and connectivity to the mother (**Wooding & Burton, 2008; Roberts *et al*., 2016**). Placentation in some mammalian species is characterized by trophoblast cells migrating into the maternal uterus where they modify the vasculature facilitating maternal nutrient flow to the placenta (**Pijnenborg *et al*., 1981; Soares *et al*., 2018**). This type of placenta is referred to as hemochorial (**Wooding & Burton, 2008; Roberts *et al*., 2016**). The human and rat possess hemochorial placentation where invasive trophoblast cells migrate deep into the uterine parenchyma (**Pijnenborg *et al*., 1981; Soares *et al*., 2018**). Regulation of deep hemochorial placentation is poorly understood. The rat represents a useful animal model for investigating the regulation of deep hemochorial placentation (**Pijnenborg & Vercruysse, 2010; Soares *et al*., 2012; Shukla & Soares, 2022**).

The rat placenta can be divided into two main compartments: i) junctional zone; ii) labyrinth zone (**Ain *et al*., 2006; Soares *et al*., 2012**). The junctional zone compartment of the placenta is situated proximal to the uterine endometrium and is responsible for transforming the maternal environment, whereas the labyrinth zone is located between the junctional zone and fetus where it facilitates nutrient delivery to the fetus (**Knipp *et al*., 1999; Soares *et al*., 2012**). Junctional zone-specific functions include the production of peptide and steroid hormones that target maternal organs and the generation of invasive trophoblast cells that migrate into and restructure the uterine parenchyma (**Soares *et al*., 1996, 2012**). The extravillous trophoblast cell column is a structure within the human placentation site, which shares some of these same responsibilities (**Soares *et al*., 2018; Knöfler *et al*., 2019**). The junctional zone and extravillous trophoblast cell column are pivotal to the regulation of maternal adaptations to pregnancy, yet little is known about how they are regulated.

In this report, we focus on the phosphatidylinositol 3-kinase (**PI3K**)/AKT pathway and its involvement in regulating junctional zone biology. AKT1 is one of three AKT serine/threonine kinases and represents an integral component of signal transduction pathways regulating cell proliferation, differentiation, migration, survival, and metabolism (**Manning & Toker, 2017; Cole *et al*., 2019**). AKT1 has also been implicated in placentation and trophoblast cell development through rodent mutagenesis experiments and investigations with human trophoblast cells (**Kamei *et al*., 2002; Yang *et al*., 2003; Qiu *et al*., 2004; Dash *et al*., 2005; Kent *et al*., 2010, 2011, 2012; Plaks *et al*., 2011; Haslinger *et al*., 2013; Sharma *et al*., 2016**). Disruptions in AKT signaling have been connected to trophoblast cell dysfunction leading to recurrent pregnancy loss, preeclampsia, and infertility (**Pollheimer & Knöfler, 2005; Ferretti *et al*., 2007; Fisher, 2015; Burton & Jauniaux, 2018**). Herein we show that AKT1 inactivation leads to placental and fetal growth restriction in the rat. AKT1 acts via phosphorylation of its target proteins leading to functional changes, including activation or inhibition of target protein function (**Manning & Toker, 2017; Cole *et al*., 2019**). We identified forkhead box O4 (**FOXO4**), a transcription factor, as an AKT1 substrate within the rat junctional zone and in rat trophoblast cells and demonstrated FOXO4 involvement in junctional zone development and the regulation of trophoblast cell differentiation.

## RESULTS

### Generation of an *Akt1* mutant rat model

AKT1 and phosphorylated AKT are ubiquitously expressed throughout the rat placentation site (**Fig. S1A**). We examined the role of AKT1 in regulating deep placentation in the rat using CRISPR/Cas9 genome editing. A mutant rat strain possessing a 1,332 bp deletion within the *Akt1* gene was generated (**Fig. 1A and B**). The deletion included part of Exon 4, the entire region spanning Exon 5 through Exon 6, and part of Exon 7 and led to a frameshift and premature stop codon (**Fig. 1A and B**). The deletion effectively removed the kinase domain and regulatory regions of AKT1 (**Fig. 1C**). The *Akt1* mutation was successfully transmitted through the germline. A rat colony possessing the *Akt1* mutation was established and maintained via heterozygous x heterozygous breeding. Mating of heterozygotes produced the predicted Mendelian ratio (**Fig. 1D; Table S1**). Placental tissues possessing a homozygous deletion within the *Akt1* locus (*Akt1^-/-^*) were deficient in AKT1 protein and exhibited prominent deficits in pan-AKT and phospho-AKT protein expression (**Fig. 1E**). Residual AKT activity could have arisen from AKT2 or AKT3, as previously demonstrated for rat trophoblast cells (**Kent et al., 2011**). These findings support the successful disruption of the *Akt1* locus and are consistent with AKT1 being the predominant AKT isoform within the placenta (**Yang *et al*., 2003; Kent *et al*., 2011; Haslinger *et al*., 2013**).

**Figure 1.**
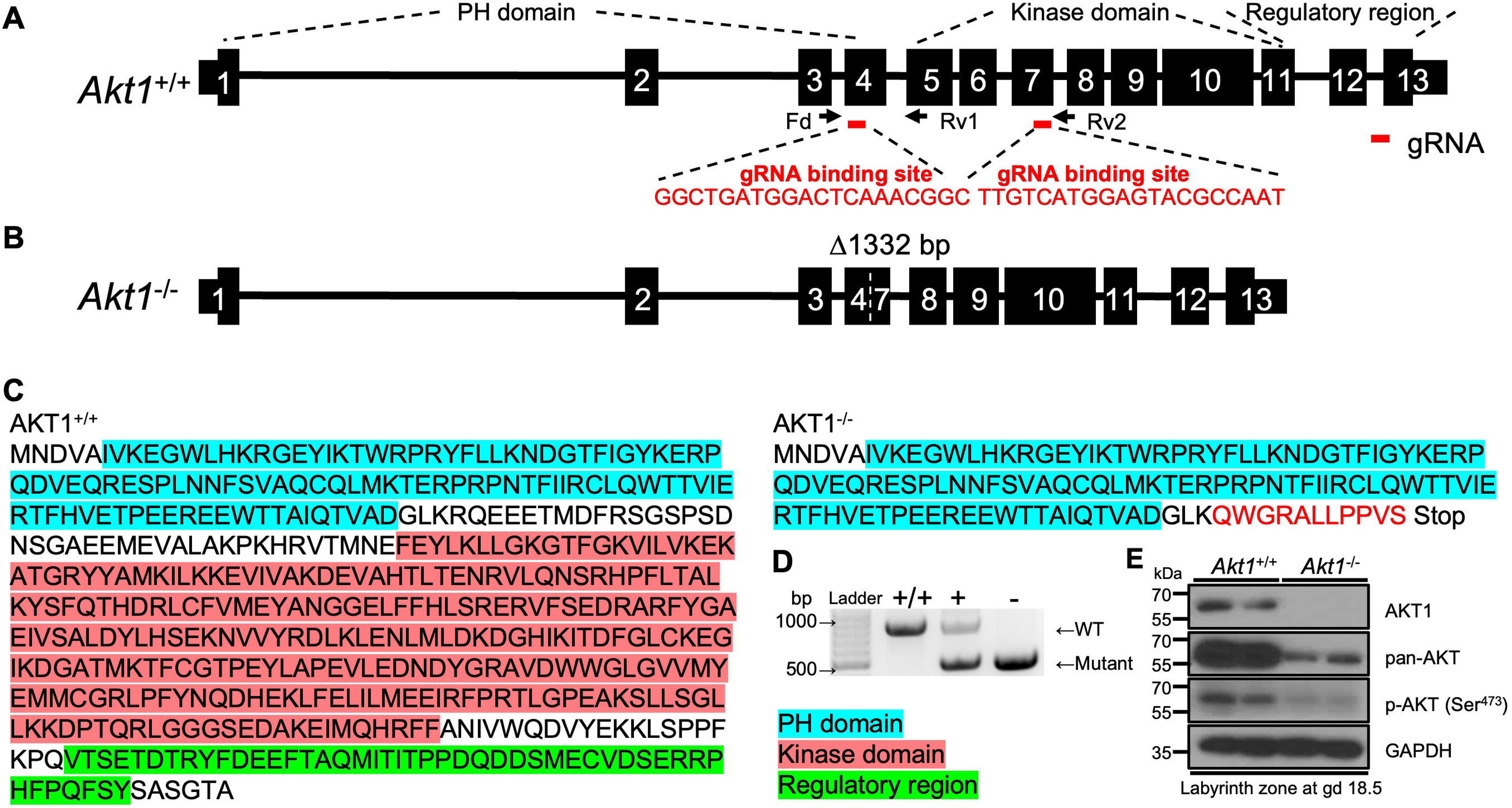
In vivo genome editing of the rat *Akt1* locus. **A)** Schematic representation of the rat *Akt1* gene (*Akt1*^+/+^) and guide RNA target sites within Exons 4 and 7 (NM_033230.3). Red bars beneath Exons 4 and 7 correspond to the 5’ and 3’ guide RNAs used in the genome editing. **B)** The mutant *Akt1* allele (*Akt1*^-/-^) possesses a 1,332 bp deletion. Parts of Exons 4 and 7 and all of Exons 5 and 6 are deleted, leading to a frameshift and premature Stop codon in Exon 7. **C)** Amino acid sequences for AKT1^+/+^ and AKT1^-/-^. The red sequence corresponds to the frameshift in Exon 7. The blue, red, and green highlighted amino acid sequence regions correspond to the pleckstrin homology (**PH**), kinase, and regulatory domains, respectively. **D)** Offspring were backcrossed to wild type rats, and heterozygous mutant rats were intercrossed to generate homozygous mutants. Wild type (+/+), heterozygous (+/-), and homozygous mutant (-/-) genotypes were detected by PCR. **E)** Western blot analysis of AKT1, pan-AKT, phospho (p)-AKT (Ser^473^) protein in *Akt1*^+/+^ and *Akt1*^-/-^ placentas at gd 18.5. GAPDH was used as a loading control.

### AKT1 deficiency results in placental, fetal, and postnatal growth restriction

Disruption of the *Akt1* locus in the mouse disrupts placental, fetal, and postnatal growth (**Chen *et al*., 2001; Cho *et al*., 2001; Yang *et al*., 2003; Plaks *et al*., 2011; Kent et al., 2012**). We observed a similar phenotype in the rat. Gestation day (**gd**) 18.5 placental and fetal weights and postnatal pup weights were significantly smaller in the *Akt1*^-/-^ rat model when compared to *Akt1*^+/+^ rats (**Fig. 2A-H**). Junctional and labyrinth zone compartments of the placenta were also significantly smaller in *Akt1*^-/-^ placentas (**Fig. 2D, E, I**).

**Figure 2.**
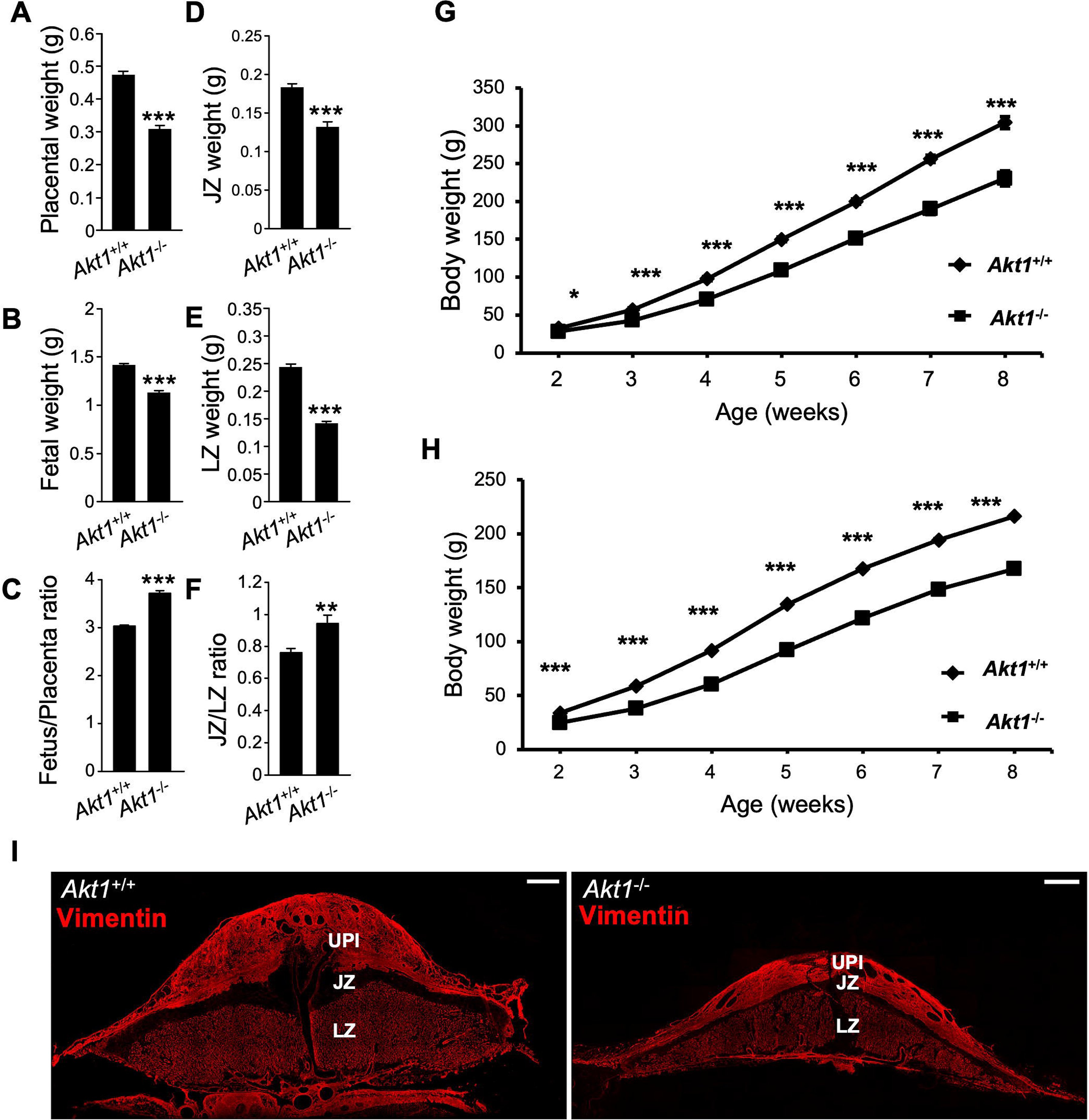
*Akt1*^-/-^ placentas and fetuses are growth restricted, and *Akt1*^-/-^ rats exhibit postnatal growth restriction. Placentas (**A**) and fetuses (**B**) were dissected from *Akt1*^+/-^ intercrosses at gd 18.5 and weighed; **C**, fetus/placenta ratio. Placentas were then separated into junctional zone (**JZ**; **D**) and labyrinth zone (**LZ**;**E**) compartments, and weighed; **F**, JZ/JZ ratio. Graphs represent means ± SEM. *Akt1*^+/+^, n = 22 from 6 dams; *Akt1*^-/-^, n = 17 from 6 dams. Asterisks denote statistical differences (\*\**P* < 0.01; \*\*\**P* < 0.001) as determined by Student’s or Welch’s *t*-test. Body weights of *Akt1*^+/+^ and *Akt1*^-/-^ pups were measured from two to eight weeks after birth: males **(G)** and females **(H)**. Graphs represent means ± SEM. n = 13-23/group. Asterisks denote statistical differences (\**P* < 0.05; \*\*\**P* < 0.001) as determined by Student’s or Welch’s *t*-test. **I**) Vimentin immunostaining of gd 18.5 *Akt1*^+/+^ and *Akt1*^-/-^ placentation sites. The junctional zone (**JZ**) is negative for vimentin immunostaining, whereas the uterine-placental interface (**UPI**) and labyrinth zone (**LZ**) stain positive for vimentin. Scale bars=1000 μm

### AKT1 regulates junctional zone and invasive trophoblast cell phenotypes

Transcript profiles were determined for *Akt1*^+/+^ and *Akt1*^-/-^ gd 18.5 junctional zone tissues using RNA-sequencing (**RNA-seq**). The size and morphological phenotypes associated with inactivation of AKT1 were associated with distinct transcript profiles (**Fig. 3; Table S2**). Disruption of AKT1 resulted in upregulation of 254 transcripts and downregulation of 333 transcripts (**Table S2**). Pathway analysis included signatures for cell cycle, DNA replication, cellular senescence, and PI3K-AKT signaling pathways (**Fig. 3A, Table S3**). Transcripts encoding cell cycle progression were consistently repressed in the *Akt1*^-/-^ junctional zones (**Fig. 3A and B**). In addition, we also observed prominent downregulation of a member of the expanded prolactin (**PRL**) gene family, *Prl8a4* (2-fold), and the upregulation of cellular communication network factor 3 (3-fold), *Ccn3*, also called *Nov*; **Fig. 3B**).

**Figure 3.**
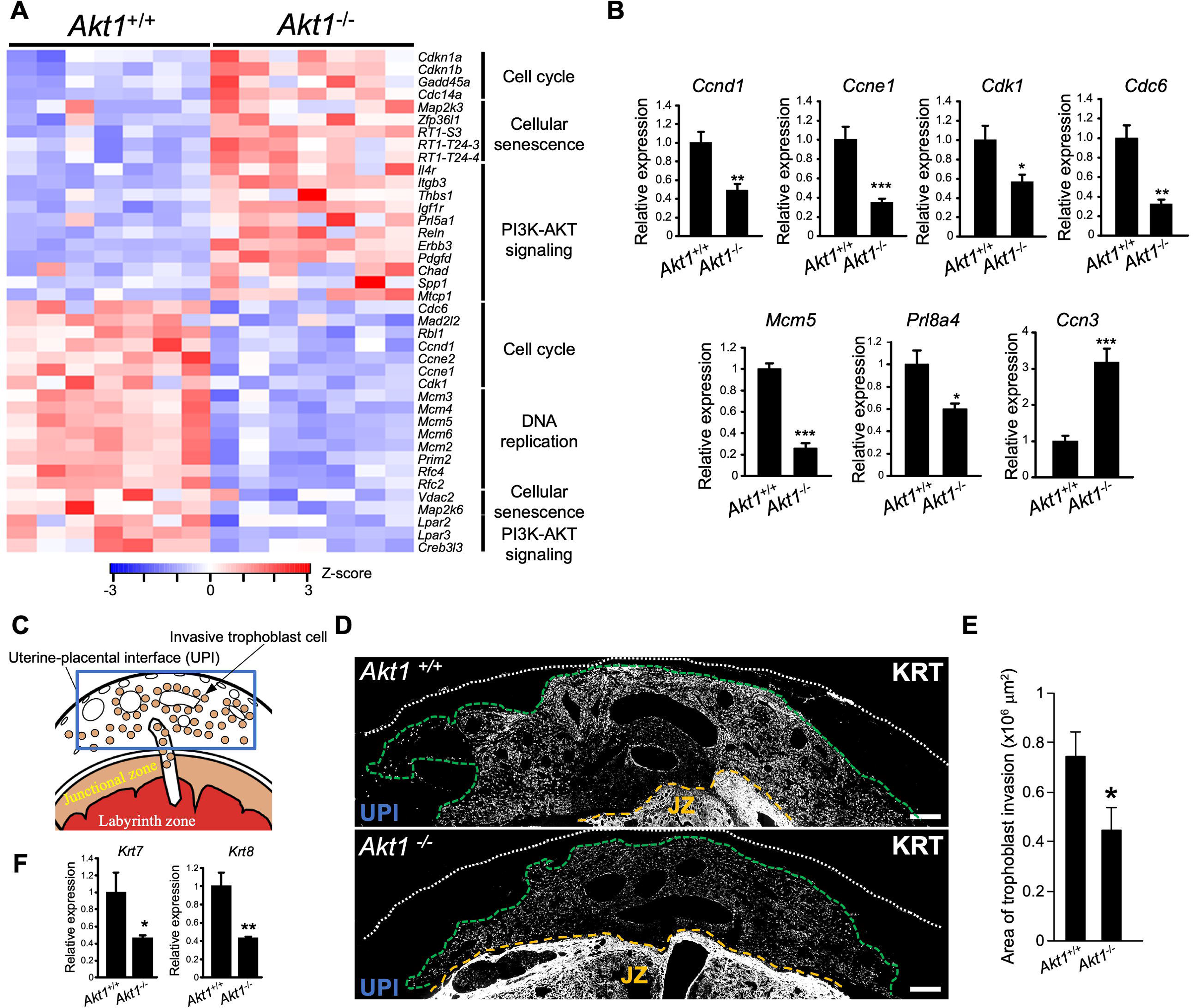
AKT1 regulates junctional zone and invasive trophoblast cell phenotypes. **A)** Heat maps depicting differentially expressed genes between *Akt1*^+/+^ and *Akt1*^-/-^ junctional zones. The heatmap color keys represent z-scores of TPM values. **B**) RT-qPCR validation of RNA-seq results (n=6/group). **C)** Schematic representation of a late gestation placentation site. The uterine-placental interface (UPI), site for intrauterine trophoblast invasion, is highlighted in the boxed area. **D)** AKT1 deficiency affected intrauterine trophoblast cell invasion. Trophoblast cells were immunostained for pan-cytokeratin (KRT). Representative images are shown. The extent of intrauterine trophoblast invasion is demarcated using a green dashed line. Scale bars =500 μm. The white dotted line represents the outer border of the uterus, and the yellow dashed line represents the uterine border with the placenta. **E)** The area of intrauterine trophoblast invasion is graphically depicted (n = 6/group). Graphs represent means ± SEM. An asterisk denotes statistical difference (\**P* < 0.05) as determined by Student’s *t*-test. F) RT-qPCR measurements of *Krt7* and *Krt8* transcripts, signature markers for invasive trophoblast cells, within dissected uterine-placental interface tissue specimens at gd 18.5 (*Akt1*^+/+^, n = 21; *Akt1*^-/-^, n = 16). Graphs represent means ± SEM. Asterisks denote statistical difference (\**P* < 0.05; \*\**P* < 0.01; \*\*\**P* < 0.001) as determined by Student’s or Welch’s *t*-test.

The junctional zone serves as the source of invasive trophoblast cells entering the uterus. Consequently, we investigated the uterine-placental interface of *Akt1*^+/+^ and *Akt1*^-/-^ placentas and monitored the surface area occupied by intrauterine invasive trophoblast cells and the expression of invasive trophoblast cell-specific transcripts. *Akt1*^-/-^ invasive trophoblast cells exhibited decreased infiltration into the uterus (**Fig. 3C-E**). The decrease in trophoblast invasion was proportional to the size of *Akt1*^+/+^ versus *Akt1*^-/-^ junctional zone compartments (**Fig. S1B**). We also observed approximately a 50% decrease in the expression of cytokeratin transcripts, which are expressed by invasive trophoblast cells within the uterine-placental interface (**Fig. 3F**).

Collectively, the data indicate that AKT1 signaling has profound effects on development of the junctional zone and the invasive trophoblast cell lineage.

### FOXO4 is a target of PI3K/AKT signaling

Forkhead box (**FOX**) transcription factors are known targets of PI3K/AKT signaling and have key roles in regulating developmental processes (**Lam *et al*., 2013; Schmitt-Ney, 2020; Herman *et al*., 2021**). We interrogated RNA-seq datasets from wild type gd 18.5 junctional zone for FOX family transcription factors. Transcripts for several FOX transcription factors were detected (transcripts per million, TPM value ≥ 1.0) (**Fig. 4A**). *Foxo4* transcripts were striking in their abundance relative to all other FOX family members. AKT1 did not significantly affect expression levels for any of the FOX family transcripts (**Table S2**). *Foxo4* transcripts were specifically localized to the junctional zone and a subset of invasive trophoblast cells (**Fig. 4B and C**). FOXO4 protein and phosphorylated FOXO4 exhibited similar tissue distributions (**Fig. S2A**). Phosphorylated FOXO4 protein was significantly diminished in *Akt1* null junctional zones (**Fig. 4D, Fig S2B**). We next explored FOXO4 in differentiated rat trophoblast stem (**TS**) cells (**Asanoma *et al*., 2011**). Rat TS cells can differentiate into trophoblast giant cells and spongiotrophoblast cells and represent an effective in vitro model for investigating junctional zone development (**Chakraborty *et al*., 2011, 2016; Asanoma *et al*., 2012; Kubota *et al*., 2015; Muto *et al*., 2021; Varberg *et al*., 2021**). *Foxo4* transcript and total and phosphorylated FOXO4 protein showed striking increases in abundance following TS cell differentiation (**Fig. 4E-F**). As previously demonstrated, AKT activity increased following trophoblast cell differentiation (**Kamei *et al*., 2002; Kent *et al*., 2010, 2011; Fig. 4G**). AKT activation was required for optimal FOXO4 phosphorylation (**Fig. 4G**). Intracellular distributions of FOXO4 and phosphorylated FOXO4 were affected by inhibition of PI3K in differentiated rat TS cells (**Fig. 4H**). Disruption of PI3K/AKT signaling shifted FOXO4 and phosphorylated FOXO4 to the nucleus (**Fig. 4H**). Inhibition of PI3K in rat TS cells did not significantly impact a measure of apoptosis (cleaved caspase 3) but did show evidence for increased autophagy (**Fig. S2C and D**). Thus, we provide support for a link between PI3K/AKT signaling and FOXO4 in trophoblast cell lineage development.

**Figure 4.**
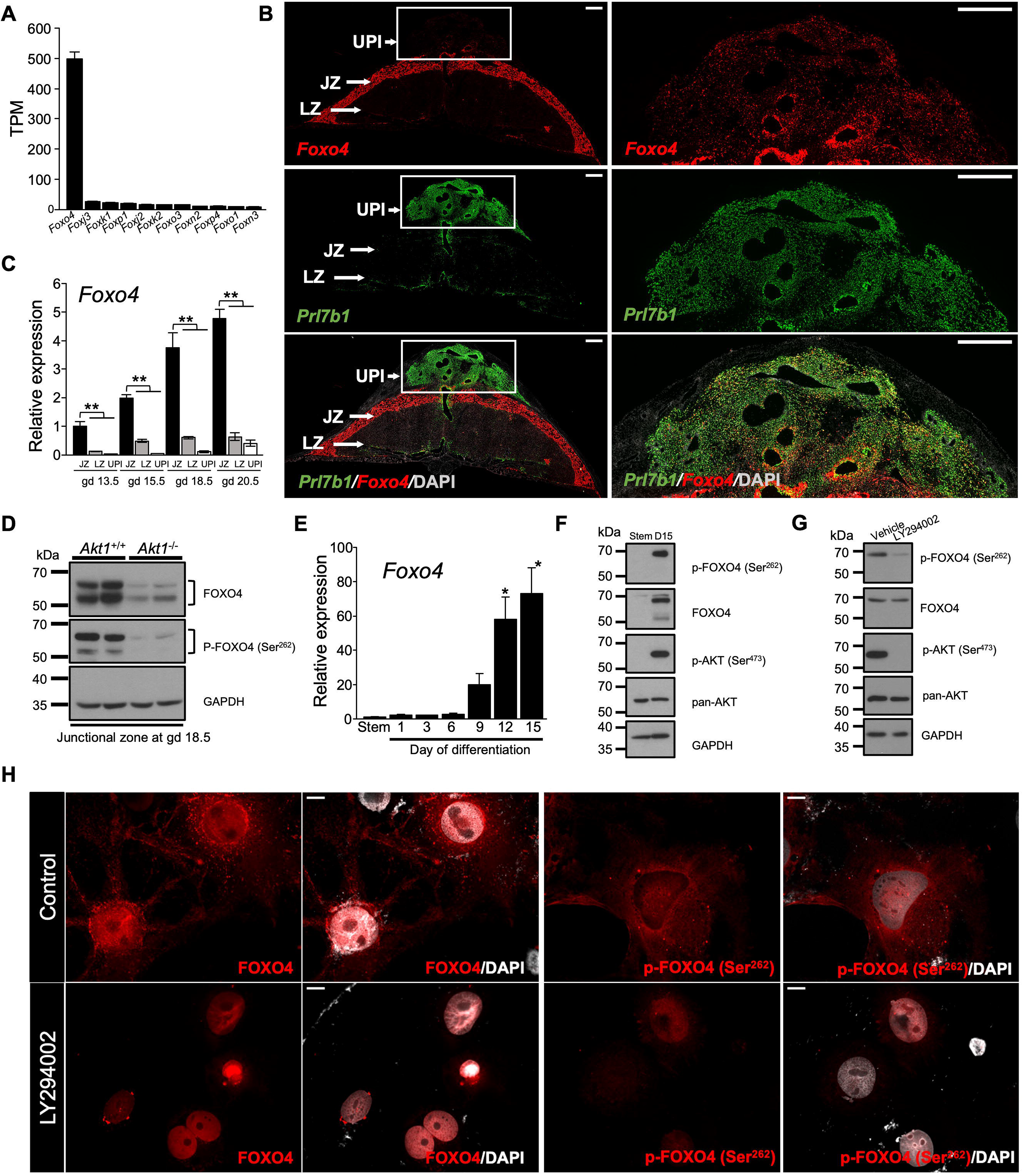
FOXO4 is a target of PI3K/AKT signaling. **A**) Expressions of transcripts for several FOX transcription factors in the junctional zone. **B**) *In situ* localization of transcripts for *Foxo4* with *Prl7b1* (invasive trophoblast-specific transcript) in the placentation site at gd 18.5 of rat pregnancy. Scale bars = 1000 μm. **C**) RT-qPCR measurements of *Foxo4* transcripts in the uterine-placental interface, junctional and labyrinth zones during the second half of gestation (n = 6-9/group). Graphs represent means ± SEM. Asterisks denote statistical difference (\*\**P* < 0.01) as determined by Steel test. UPI: uterine-placental interface; JZ: junctional zone; and LZ: labyrinth zone. **D**) Western blot analysis of phospho (p)-FOXO4 (Ser^262^) and FOXO4 proteins in *Akt1*^+/+^ and *Akt1*^-/-^ placentas at gd 18.5. GAPDH was used as a loading control. **E)** RT-qPCR measurements of *Foxo4* transcripts in the stem state and following induction of differentiation (n = 4-6/group). Graphs represent means ± SEM. Asterisks denote statistical difference (vs Stem, \**P* < 0.05) as determined by Dunnett’s test. Western blot analysis of phospho (p)-FOXO4 (Ser^262^), FOXO4, p-AKT (Ser^473^), and pan-AKT proteins in the stem and differentiating (day 15 of differentiation, D15) states **(F)**, and in the differentiating state (D15) following treated with vehicle (0.1% DMSO) or a phosphatidylinositol 3-kinase (**PI3K**) inhibitor (LY294002, 10 μM) for 1 h **(G)**. GAPDH was used as a loading control. **H**) Rat TS following 15 days in differentiating conditions were treated with vehicle (0.1% DMSO, Control) or a phosphatidylinositol 3-kinase (**PI3K**) inhibitor (LY294002, 10 μM) for 24 h and then immunostained for phospho (p)-FOXO4 (Ser^262^), FOXO4. Representative images are shown. Scale bars =50 μm.

### Generation of a *Foxo4* mutant rat model

We examined the role of FOXO4 in regulating placentation in the rat using CRISPR/Cas9 genome editing. The *Foxo4* gene consists of four exons and resides on the X chromosome (**Liu *et al*., 2020**). A mutant rat strain possessing a 3,096 bp deletion within the *Foxo4* gene was generated (**Fig. 5A and B**). The deletion included the 3’ part of Exon 2 and the 5’ part of Exon 3 and led to a frameshift and a premature stop codon (**Fig. 5A and B**). The deletion effectively disrupted the conserved forkhead DNA binding domain and removed nuclear localization, nuclear export, and transactivation domains of FOXO4 (**Fig. 5C**). The *Foxo4* mutation was successfully transmitted through the germline (**Fig. 5D; Table S4**). A rat colony possessing the *Foxo4* mutation was established and maintained via hemizygous male x wild type female breeding, which produced the predicted Mendelian ratio (**Table S4**). Junctional zone tissues possessing a maternally inherited *Foxo4* mutation (*Foxo4*^Xm-^) were deficient in FOXO4 protein (**Fig. 5E**). The results are consistent with paternal silencing of X chromosome-linked genes expressed in extraembryonic tissues (**Takagi & Sasaki, 1975; West et al., 1978; Hemberger, 2002**). FOXO4 was successfully disrupted in the rat.

**Figure 5.**
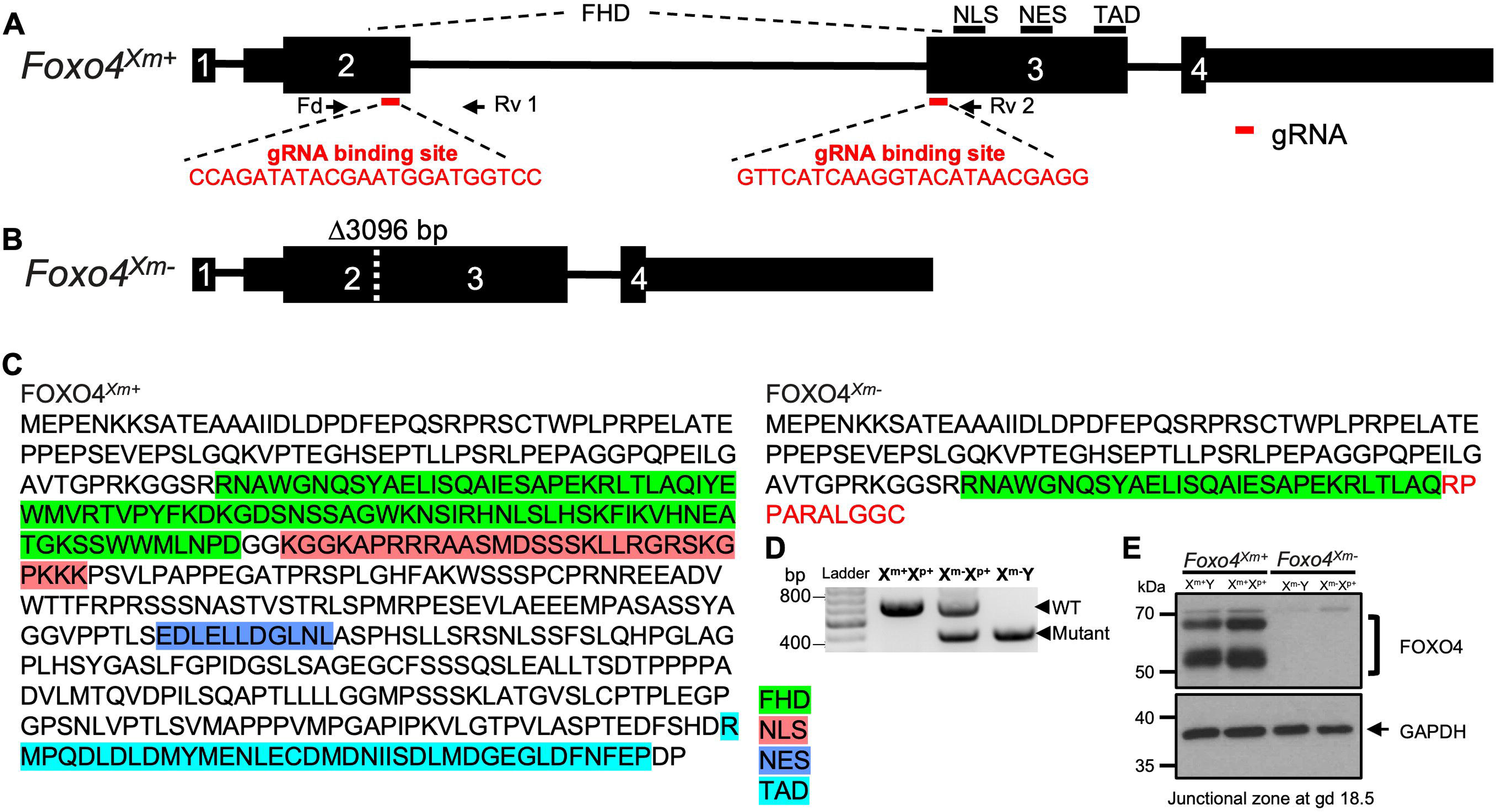
In vivo genome editing of the rat *Foxo4* locus. **A)** Schematic representation of the rat *Foxo4* gene (*Foxo4*^Xm+^) and guide RNA target sites within Exons 2 and 3 (NM_001106943.1). Red bars beneath Exons 2 and 3 correspond to the 5’ and 3’ guide RNAs used in the genome editing. **B)** The mutant *Foxo4* allele (*Foxo4*^Xm-^) possesses a 3,096 bp deletion. Parts of Exons 2 and 3 are deleted, leading to a frameshift and premature Stop codon in Exon 3. **C)** Amino acid sequences for FOXO4^Xm+^ and FOXO4^Xm-^. The red sequence corresponds to the frameshift in Exon 3. The green, red, dark blue, and light blue highlighted amino acid sequence regions correspond to the forkhead winged-helix DNA-binding domain (**FHD**), nuclear localization sequence (**NLS**), nuclear export sequence (**NES**), and transactivation domain (**TAD**), respectively. **D)** Heterozygous mutant female rats were crossed to wild type male rats to generate hemizygous null male rats. Wild type (+/+), heterozygous (+/-), and hemizygous null (-/y) genotypes were detected by PCR. **E)** Western blot analysis of FOXO4 protein in the junctional zone of *Foxo4*^Xm+^ (X^m+^Y and X^m+^X^p+^) and *Foxo4*^Xm-^ (X^m-^Y and X^m-^X^p+^) placentas at gd 18.5. GAPDH was used as a loading control.

### FOXO4 deficiency results in placentomegaly and a modified junctional zone phenotype

Placentation site phenotypes of mice possessing mutations at the *Foxo4* locus have not been described (**Liu *et al*., 2020; Hosaka *et al*., 2004**). Rats possessing a maternally inherited mutant *Foxo4* allele (*Foxo4*^Xm-^) exhibited placentomegaly and decreased placental efficiency (fetal/placental weight ratio) when examined on gd 18.5 (**Fig. 6A-C**) and gd 20.5 (**Fig S3A-C**). In contrast, a paternally inherited mutant *Foxo4* allele did not significantly affect placenta or fetal weights (**Fig S3D and E**). FOXO4 deficiency associated placentomegaly included significantly larger junctional and labyrinth zones (**Fig. 6D-G, Fig S3F-H**) but did not affect the intrauterine invasive trophoblast cell lineage (**Fig S4**). Transcript profiles were determined for wild type (*Foxo4*^Xm+^) and *Foxo4^Xm-^* gd 18.5 junctional zone tissues using RNA-seq (**Fig. 6H**). Disruption of FOXO4 resulted in upregulation of 369 transcripts and downregulation of 845 transcripts (**Table S5**). Pathway analysis included signatures for PI3K-AKT signaling, cell cycle, DNA replication, extracellular matrix receptor interaction, and complement and coagulation pathways (**Fig. 6H, Table S6**). Cell cycle and DNA replication signatures may be contributing to the size disparities in wild type (*Foxo4*^Xm+^) versus *Foxo4^Xm-^* junctional zone compartments. Junctional adhesion molecule-like (**JAML**), lipoprotein(a) like 2 (**LPAL2**), erb-b2 receptor tyrosine kinase 3 (**ERBB3**), and growth factor receptor bound protein 7 (**GRB7**) were each conspicuous in their prominent downregulation (>90% decrease) in FOXO4 deficient junctional zone tissue (**Fig. 6I**). JAML contributes to epithelial barrier function, modulates immune cell trafficking, and angiogenesis (**Kummer & Ebnet, 2018**), whereas LPAL2 is a long noncoding RNA contributing to inflammatory and oxidative stress responses (**Han *et al*., 2018**). JAML and LPAL2 have not previously been linked to trophoblast or placental biology. ERBB3 is a receptor for neuregulin 1 and promotes trophoblast cell survival (**Fock *et al*., 2015**) and GRB7 is an adaptor protein participating in signal transduction activated through ERBB3 (**Fiddes *et al*., 1998**). Interestingly, numerous junctional zone transcripts regulated by FOXO4 were reciprocally regulated by AKT1 (88% of transcripts upregulated in *Akt1* null were downregulated in *Foxo4* mutant tissues; 38% of transcripts downregulated in *Akt1* null were upregulated in *Foxo4* mutant tissues; **Fig. 7**). The reciprocal relationship between AKT1 and FOXO4 is evident at structural and molecular levels.

**Figure 6.**
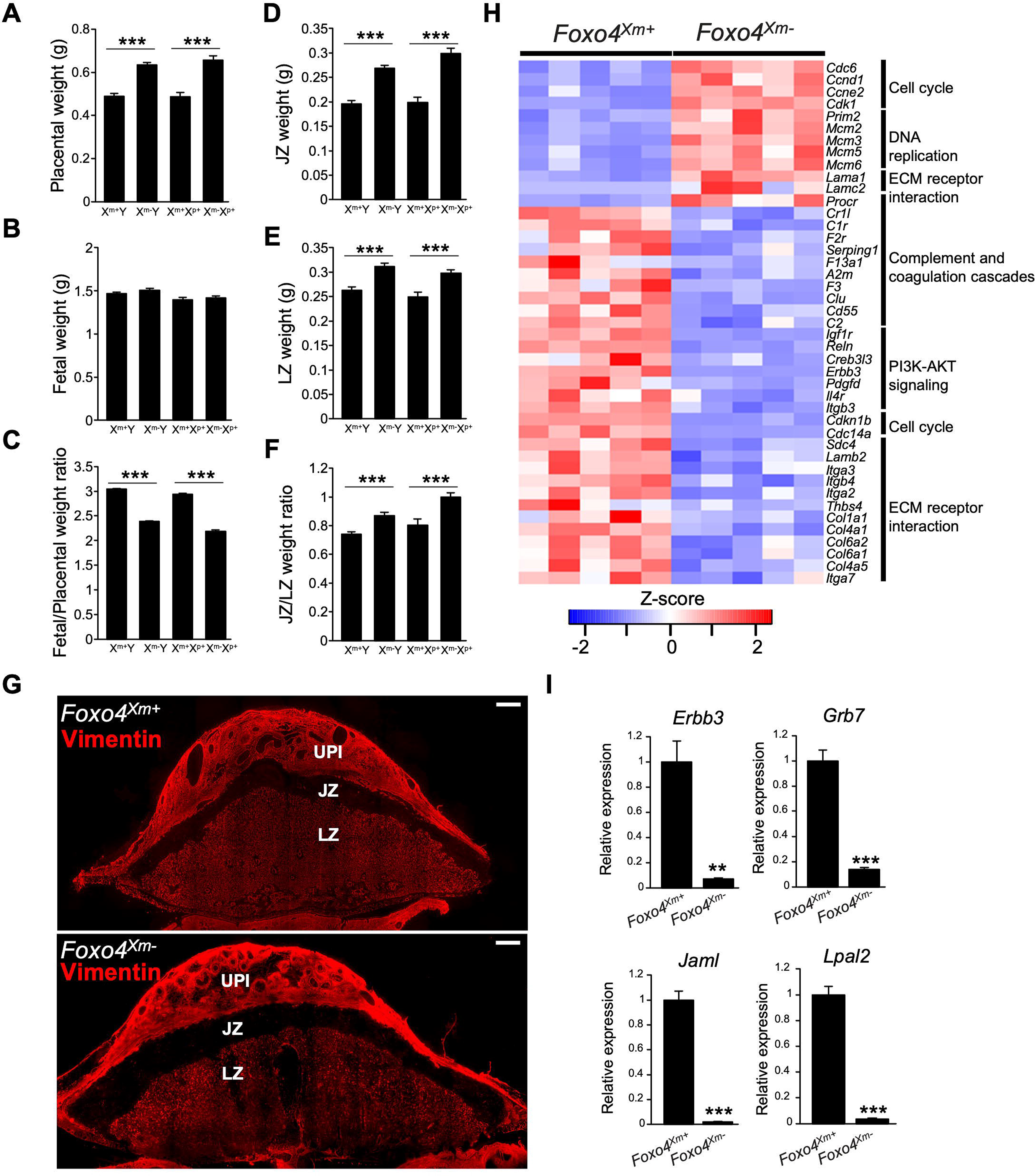
*Foxo4* hemizygous null and *Foxo4* maternally inherited heterozygous conceptuses exhibit placental overgrowth, and FOXO4 deficiency alters the transcriptomes of the junctional zone. Placentas **(A)** and fetuses **(B)** were dissected from *Foxo4* heterozygous females mated with wild type males at gd 18.5 and weighed; **C**) fetus/placenta ratio. Placentas were then separated into junctional zone (**JZ, D**) and labyrinth zone (**LZ, E**) compartments, and weighed; **F**, JZ/LZ weight ratio. Graphs represent means ± SEM. X^m+^Y, n = 25; X^m-^Y, n = 31; X^m+^X^p+^, n = 14; X^m-^X^p+^, n = 22 from 8 dams. Asterisks denote statistical differences (\*\*\**P* < 0.001) as determined by Student’s or Welch’s *t*-test. **G**) Vimentin immunostaining of gd 18.5 *Foxo4*^Xm+^ and *Foxo4*^Xm^ placentation sites. The junctional zone (**JZ**) is negative for vimentin immunostaining, whereas the uterine-placental interface (**UPI**) and labyrinth zone (**LZ**) stain positive for vimentin. Scale bars=1000 μm. **H**) Heat maps depicting differentially expressed genes in *Foxo4*^Xm+^ versus *Foxo4*^Xm-^ junctional zones. The heatmap color keys represent z-scores of TPM values. **I**) RT-qPCR validation of RNA-seq results (n=6/group). Graphs represent means ± SEM. Asterisks denote statistical difference (**P<0.01; ***P<0.001) as determined by Student’s or Welch’s t-test.

**Figure 7.**
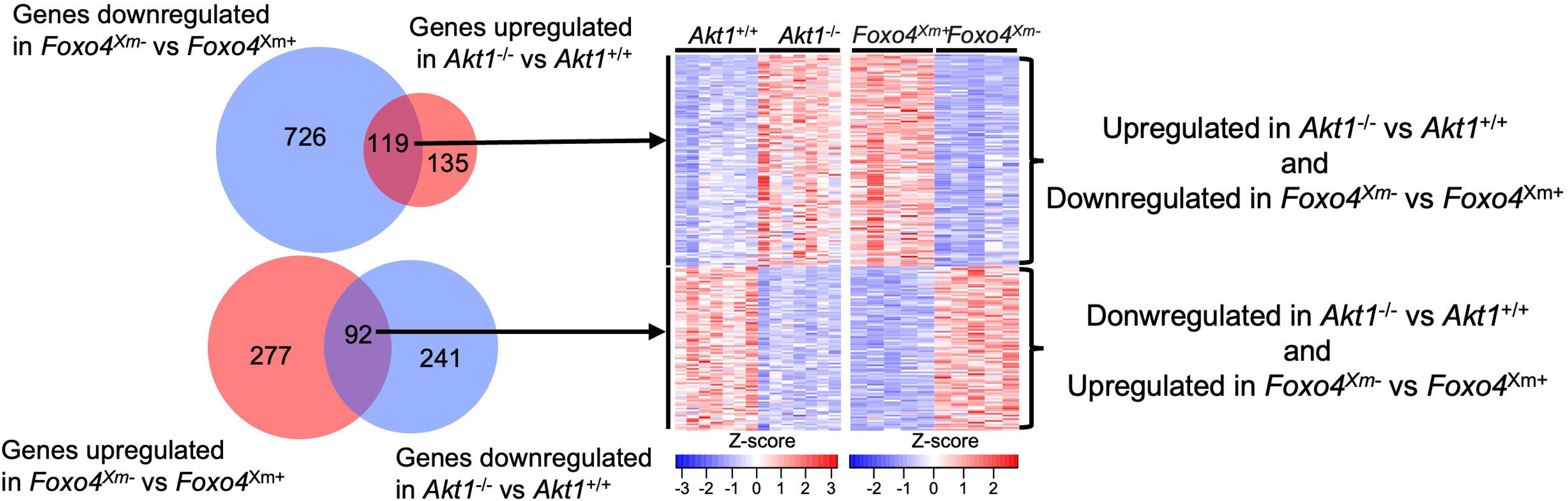
Reciprocal relationship between AKT1 and FOXO4 in the junctional zone. Venn diagram and heatmaps representing overlap of differentially expressed genes inversely regulated by AKT1 and FOXO4. The heatmap color keys represent z-scores of TPM values.

### FOXO4 contributes to the regulation of the trophoblast cell lineage

We next modeled junctional zone cell biology using rat TS cells. The consequences of FOXO4 disruption in differentiating rat TS cells were examined. FOXO4 expression was inhibited via ectopic expression of short hairpin RNAs specific to *Foxo4* (**Fig. 8A and B**). Although, a morphologic phenotype was not evident, prominent differences in the transcriptomes of TS cells expressing control versus *Foxo4* shRNAs were observed (**Fig. 8C**). Disruption of FOXO4 resulted in upregulation of 260 transcripts and downregulation of 443 transcripts (**Table S7**). Pathway analysis included signatures for PI3K-AKT signaling, longevity regulating, calcium signaling, and glutathione metabolism pathways (**Fig. 8C, Table S8**). Among the dysregulated transcripts was an upregulation of matrix metallopeptidase 12 (2-fold), a known constituent of endovascular invasive trophoblast cells (**Harris et al., 2010; Chakraborty et al., 2016**) and downregulation of trophoblast specific protein alpha (3-fold), a transcript characteristic of spongiotrophoblast cells within the junctional zone (**Iwatsuki *et al*., 2000**). FOXO4 is a known regulator of responses to oxidative stress (**Liu *et al*., 2020**). Several transcripts associated with inflammatory and cellular stress responses, including thioredoxin interacting protein (7-fold), glutathoione S-transferase mu 1 (3-fold), arachidonate 5-lipoxygenase activating protein (2-fold), interferon kappa (10-fold), nuclear protein 1 (2.7-fold), and *Lpal2* (2.2-fold), were prominently downregulated (**Sies & Cadenas, 1985; LaFleur *et al*., 2001; Mashima & Okuyama, 2015; Han *et al*., 2018; Huang *et al*., 2021; Nirgude & Choudhary, 2021; Qayyum *et al*., 2021; Satapathy & Wilson, 2021**).

**Figure 8.**
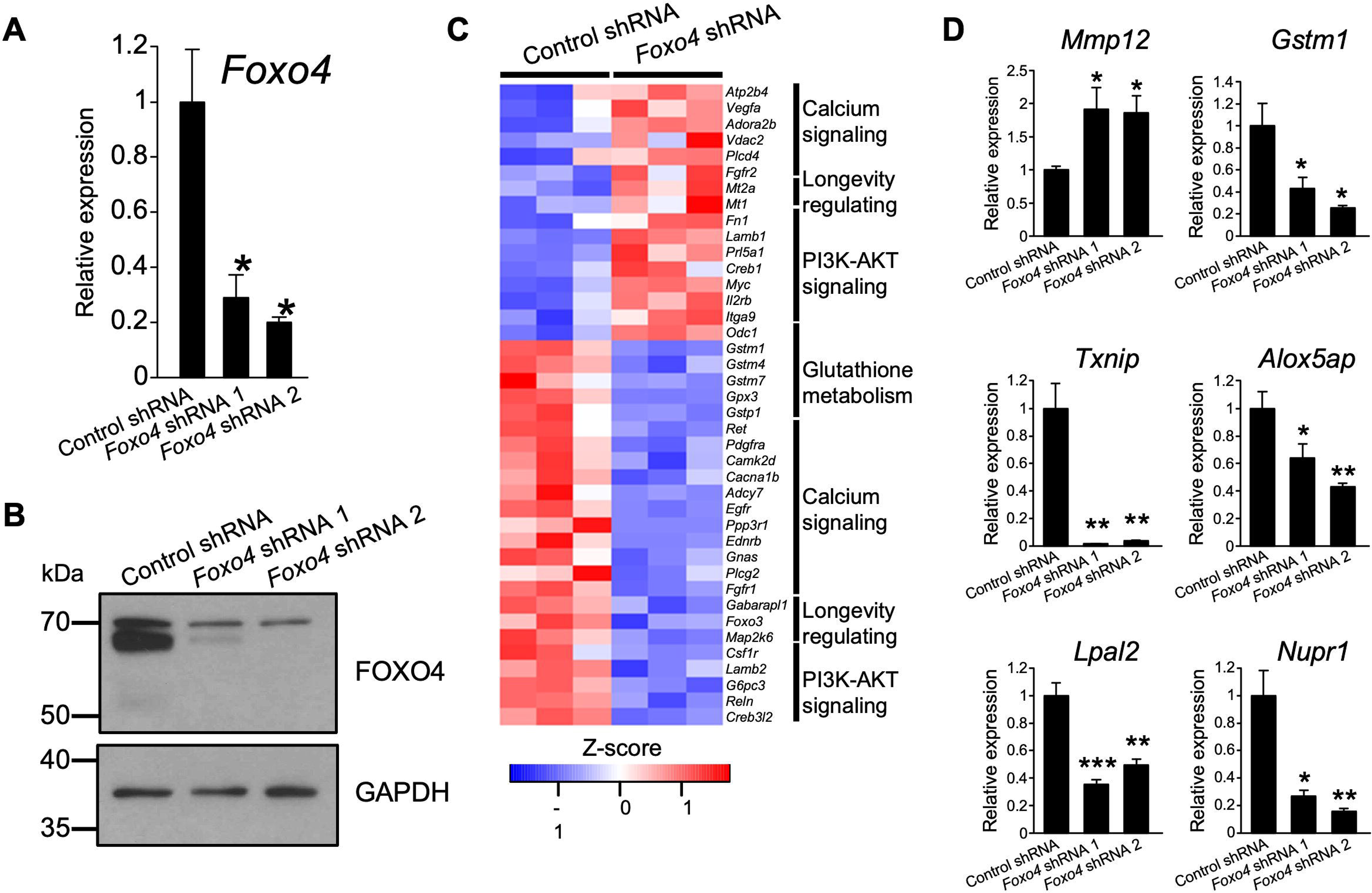
FOXO4 alters the rat TS cell. FOXO4 knockdown efficiency was validated by RT-qPCR (**A**, n = 4/group) or western blot **(B)** analyses in rat TS cells at day 15 of differentiation following transduction with lentivirus containing a control shRNA or one of two independent *Foxo4-specific* shRNAs. Graphs represent means ± SEM. Asterisks denote statistical difference (vs Control shRNA, \**P* < 0.05) as determined by Student’s or Welch’s *t*-test. **C)** Heatmap depicting differentially expressed genes between control and *Foxo4* shRNA-treated rat TS cells. **D)** RT-qPCR validation of RNA-seq results (Control shRNA, n = 4; *Foxo4* shRNA 1, n = 4; *Foxo4* shRNA 2, n = 4). Graphs represent means ± SEM. Asterisks denote statistical difference (compared to Control shRNA, \**P* < 0.05; \*\**P* < 0.01; \*\*\**P* < 0.001) as determined by Student’s or Welch’s *t*-test.

In vivo and in vitro disruptions of FOXO4 were consequential but did not yield identical transcriptomic effects. These discrepancies may reflect intrinsic differences in the cellular compositions of the gd 18.5 junctional zone versus differentiated rat TS cells, in vivo versus in vitro cell environments, or the null in vivo model versus the hypomorphic in vitro model.

### Key findings

Collectively, the results indicate that AKT1 drives placental growth, including regulation of deep intrauterine trophoblast cell invasion. These actions are accomplished, at least in part, through modulation of FOXO4, which acts to restrain placental growth and coordinate responses to physiological stressors.

## DISCUSSION

The rat and human possess a type of hemochorial placentation where specialized trophoblast cells penetrate deep into the uterus and transform the uterine parenchyma, including the vasculature (**Pijnenborg *et al*., 1981; Pijnenborg & Vercruysse, 2010; Soares *et al*., 2012**). Central to deep placentation is the source of invasive trophoblast cells, which in the rat is a compartment within the placenta referred as the junctional zone and, in the human, the extravillous trophoblast cell column (**Soares *et al*., 2012, 2018; Knöfler *et al*., 2019**). In addition to an intrauterine role, the cellular constituents of these placental compartments produce hormones directed to maternal tissues with actions that ensure in utero survival and promotion of fetal growth (**Soares *et al*., 1996; John, 2017**). In this report, AKT1 and FOXO4 were identified as regulators of rat junctional zone development. In vivo disruption of AKT1 and FOXO4 led to opposite effects on junctional zone development. AKT1 deficiency resulted in growth restriction of the junctional zone, phenotypic alteration of the invasive trophoblast cell lineage, as well as compromised fetal and postnatal growth, and AKT1 was capable of phosphorylating FOXO4 in rat trophoblast cells. Deficiency of FOXO4 resulted in an expanded junctional zone. Deficits in AKT1 or FOXO4 also impacted transcriptomic profiles of the junctional zone. The findings indicate that AKT1 and FOXO4 are part of a gene regulatory network controlling hemochorial placentation.

AKT1 signaling influenced placental development. *Akt1* null mutations in the mouse and rat yield similar phenotypes characterized by placental, fetal, and postnatal growth restriction (**Chen *et al*., 2001; Cho *et al*., 2001; Yang *et al*., 2003; Plaks *et al*., 2011; Kent et al., 2012**). Smaller junctional zones accompanying AKT1 deficiency were associated with a downregulation of transcripts encoding proteins driving cell proliferation. These results imply that size differences of junctional zone compartments in the wild type versus *Akt1* nulls were related to, at least in part, diminished trophoblast cell proliferation in *Akt1* junctional zone tissues. The data also fit well with known actions of AKT signaling promoting cell proliferation in a wide range of cell types (**Manning & Cantley, 2007; Manning & Toker, 2017; Cole *et al*., 2019**). AKT1 disruption also altered differentiated junctional zone trophoblast cell phenotypes. As cellular constituents of the junctional zone differentiate, they acquire the capacity to express several members of the expanded prolactin (**PRL**) family of hormones/cytokines (**Soares, 2004; Alam *et al*., 2006; Soares *et al*., 2007**). The expression of PRL8A4, a member of the expanded PRL family, was dramatically downregulated in *Akt1* null junctional zones. PRL8A4 is an orphan ligand with little known of its significance to the biology of pregnancy other than as a signature feature of the differentiated junctional zone phenotype (**Iwatsuki *et al*., 1998; Soares *et al*., 2007**). AKT signaling has previously been implicated in the regulation of the differentiation of rodent and human trophoblast cells (**Kamei *et al*., 2003; Kent *et al*., 2010, 2011; Haslinger *et al*., 2013**). Disruption of *Akt1* also interfered with invasive trophoblast cell development. Trophoblast cell infiltration into the uterine-placental interface was diminished in *Akt1* nulls. Involvement of AKT signaling has also been implicated in the development of the human extravillous trophoblast cell lineage (**Pollheimer & Knöfler, 2005; Haslinger *et al*., 2013; Morey *et al*., 2021**). AKT signaling could affect invasive trophoblast cell development through its actions on their origin in the junctional zone and EVT cell column or alternatively, their maturation as they invade into the uterus. Finally, the impact of AKT signaling in invasive trophoblast cell development may be more profound than observed with AKT1 deficiency due to compensatory activities of AKT2 and AKT3 (**Kent *et al*., 2011; Haslinger *et al*., 2013**).

AKT1 regulates cellular function through its actions as a serine/threonine kinase and thus, phosphorylation of its substrates (**Manning & Cantley, 2007; Manning & Toker, 2017; Cole *et al*., 2019**). The forkhead box (**FOX**) family of transcription factors are well-established targets of AKT action (**Lam *et al*., 2013; Schmitt-Ney, 2020; Herman *et al*., 2021**). Among FOX family members, FOXO4 expression was uniquely elevated in the rat junctional zone. FOXO4 phosphorylation state in trophoblast cells was affected by AKT signaling. AKT-mediated phosphorylation of FOXO4 leads to FOXO4 exit from the nucleus and inactivation (**Schmitt-Ney, 2020; Herman *et al*., 2021**), which might suggest that an AKT1 deficiency would result in the stabilization of FOXO4 protein in the junctional zone. Instead, AKT1 deficiency led to depletion of junctional zone total and phosphorylated FOXO4 proteins. Consequently, the observed *Akt1* null placental phenotype was associated with the depletion of both AKT1 and FOXO4 proteins. In addition to regulation by AKT signaling, FOXO4 activity/stability is stimulated by Jun kinase and monoubiquitylation (**Essers *et al*., 2004; van der Horst *et al*., 2006; Brenkman *et al*., 2008; Liu *et al*., 2020**), while inhibited by acetylation and polyubiquitylation (**Fukuoka *et al*., 2003; Huang & Tindall, 2011; Liu *et al*., 2020**). Whether AKT1 indirectly affects FOXO4 protein via impacting these other FOXO4 regulators remains to be determined.

FOXO4 is a transcription factor implicated in the regulation of the cell cycle, apoptosis, responses to oxidative stress, and a range of disease processes (**Liu *et al*., 2020**). The original characterization of the *Foxo4* null mouse concluded that FOXO4 did not have a singular role in the pathophysiology of the mouse (**Hosaka *et al*., 2004**). The absence of a reported phenotype for the *Foxo4* null mouse model was attributed to the compensatory actions of other members of the FOXO family, including FOXO1, FOXO3, and possibly FOXO6 (**Hosaka *et al*., 2004; Liu *et al*., 2020**). We describe a prominent placental phenotype for the *Foxo4* null rat model. The placental anomalies associated with FOXO4 deficiency were compatible with the production of viable offspring. The absence of a fertility defect in the *Foxo4* null mouse model likely precluded a closer examination of placentation (**Hosaka *et al*., 2004**). However, it is also possible that elements of FOXO4 action are species specific. FOXO4 is prominently expressed in the junctional zone and to a lesser extent in invasive trophoblast cells. Disruption of FOXO4 led to an expansion of both junctional and labyrinth zone placental compartments, which probably reflects cell autonomous and non-cell autonomous actions, respectively. A striking reciprocal pattern of AKT1 versus FOXO4 gene regulation within the junctional zone was demonstrated and included differentially regulated transcripts encoding proteins involved in the regulation of cell proliferation and cell death. We surmise that AKT1 promotes junctional zone growth via stimulating the expression of transcripts connected to cell cycle progression and inhibited transcripts connected to cell death, whereas the converse is true for FOXO4. These biological roles are consistent with the known actions of AKT1 and FOXO4 in other cell systems (**Manning & Toker, 2017; Liu *et al*., 2020; Herman *et al*., 2021**). A transcriptional regulatory network involving FOXO4 has also been identified in human extravillous trophoblast cells (**Morey *et al*., 2021**). Importantly, FOXO4 also regulates trophoblast cell responses to oxidative stress and is thus, positioned to contribute to placental adaptations to a compromised maternal environment and in disease states affecting placentation.

## MATERIALS AND METHODS

### Animals

Holtzman Sprague-Dawley rats were maintained in an environmentally controlled facility with lights on from 0600 to 2000 h with food and water available ad libitum. Time-mated pregnancies were established by co-housing adult female rats (8-10 weeks of age) with adult male rats (>10 weeks of age). Detection of sperm or a seminal plug in the vagina was designated gd 0.5. Pseudopregnant females were generated by co-housing adult female rats (8-10 weeks of age) with adult male vasectomized males (>10 weeks of age). The detection of seminal plugs was designated pseudopregnancy day 0.5. Four to five-week-old donor rats were superovulated by intraperitoneal injection of pregnant mare serum gonadotropin (30 units, G4877, Sigma-Aldrich, St. Louis, MO), followed by an intraperitoneal injection of human chorionic gonadotropin (30 units, C1063, Sigma-Aldrich) ~46 h later, and immediately mated with adult males. Zygotes were flushed from oviducts the next morning (gd 0.5). The University of Kansas Medical Center (**KUMC**) Animal Care and Use Committee approved all protocols involving the use of rats.

### Tissue collection and analysis

Rats were euthanized by CO_2_ asphyxiation at designated days of gestation. Uterine segments containing placentation sites were frozen in dry ice-cooled heptane and stored at −80°C until processed for histological analyses. Alternatively, placentation sites were dissected into placentas, the adjacent uterine-placental interface tissue (also referred to as the metrial gland), and fetuses as previously described (**Ain *et al*., 2006**). Placentas were weighed and dissected into junctional zone and labyrinth zone compartments (**Ain *et al*., 2006**). Placental compartments and uterine-placental interfaces were frozen in liquid nitrogen and stored at −80°C until used for biochemical analyses. Fetuses were weighed, genotyped, and sex determined by polymerase chain reaction (**PCR**) (**Dhakal & Soares, 2017**).

### Generation of *Akt1* and *Foxo4* mutant rat models

Mutations at *Akt1* and *Foxo4* loci were generated using CRISPR/Cas9 genome editing (**Kaneko, 2017; Iqbal *et al*., 2021**). Guide RNAs targeting Exon 4 (target sequence: GCCGTTTGAGTCCATCAGCC; nucleotides 356-375) and Exon 7 (target sequence: TTGTCATGGAGTACGCCAAT; nucleotides 712-731) of the *Akt1* gene (NM_033230.3) or targeting Exon 2 (target sequence: CCAGATATACGAATGGATGGTCC; nucleotides 517-539) and Exon 3 (target sequence: GTTCATCAAGGTACATAACGAGG; nucleotides 631-653) of the *Foxo4* gene (NM_001106943.1) were electroporated into single-cell rat embryos using the NEPA21 electroporator (Nepa Gene Co Ltd, Ichikawa City, Japan). Electroporated embryos were transferred to oviducts of day 0.5 pseudopregnant rats. Initially, offspring were screened for *Akt1* or *Foxo4* mutations from genomic DNA from tail-tip biopsies using the REDExtract-N-Amp™ Tissue PCR kit (XNAT, Millipore Sigma, Burlington, MA). PCR was performed on the purified DNA samples using primers flanking the guide RNA sites, and products resolved by agarose gel electrophoresis and ethidium bromide staining. Genomic DNA containing potential mutations was amplified by PCR, gel purified, and precise boundaries of deletions determined by DNA sequencing (Genewiz Inc., South Plainfield, NJ). Founders with *Akt1* or *Foxo4* mutations were backcrossed to wild type rats to demonstrate germline transmission. Routine genotyping was performed by PCR on genomic DNA with specific sets of primers (**Table S9**).

### Western blot analysis

Tissue lysates were prepared with radioimmunoprecipitation assay lysis buffer system (sc-24948A, Santa Cruz Biotechnology, Santa Cruz, CA). Protein concentrations were determined using the *DC*™ Protein Assay Kit (5000112, Bio-Rad Laboratories, Hercules, CA). Proteins (20 μg/lane) were separated by SDS-PAGE. Separated proteins were electrophoretically transferred to polyvinylidene difluoride membranes (10600023, GE Healthcare, Milwaukee, WI) for 1 h at 100 V on ice. Membranes were subsequently blocked with 5% milk or 5% bovine serum albumin for 1 h at room temperature and probed separately with specific primary antibodies to AKT1 (1:1,000 dilution, 75692, Cell Signaling Technology, Danvers, MA), pan-AKT (1:1,000 dilution, 4691, Cell Signaling Technology), phospho-Ser^473^ AKT (1:2,000 dilution, 4060, Cell Signaling Technology), FOXO4 (1:2,000 dilution, 21535-1-AP, Proteintech, Rosemont, IL), phospho-Ser^262^ FOXO4 (1:3,000 dilution, ab126594, Abcam, Cambridge, MA), cleaved caspase-3 (Asp175, 1:1,000 dilution, 9664, Cell Signaling Technology), serine palmitoyltransferase long chain base subunit 3 (**LC3B**, 1:1,000 dilution, 2775, Cell Signaling Technology), and glyceraldehyde 3-phosphate dehydrogenase (**GAPDH**, 1:5,000 dilution, ab8245, Abcam) in Tris-buffered saline with Tween 20 (**TBST**) overnight at 4□. After primary antibody incubation, the membranes were washed in TBST three times for ten min each at room temperature. After washing, the membranes were incubated with anti-rabbit or anti-mouse immunoglobulin G (**IgG**) conjugated to horseradish peroxidase [**HRP**, 1:5,000 dilution or 1:20,000 dilution (phospho-Ser^262^ FOXO4), 7074S and 7076S, Cell Signaling Technology] in TBST for 1 h at room temperature, washed in TBST three times for ten min each at room temperature, immersed in Immobilon Crescendo Western HRP Substrate (WBLUR0500, Sigma-Aldrich), and luminescence detected using Radiomat LS film (Agfa Healthcare, Mortsel, Belgium) or Chemi Doc MP Imager (Bio-Rad)

### Transcript analysis

Total RNA was extracted from tissues using TRI Reagent Solution (AM9738, Thermo-Fisher, Waltham, MA) according to the manufacturer’s instructions. Total RNA (1 μg) was reverse transcribed using a High-Capacity cDNA Reverse Transcription Kit (4368813, Thermo-Fisher). Complementary DNAs were diluted 1:10 and subjected to reverse transcription-quantitative PCR (**RT-qPCR**) using PowerUp SYBR Green Master Mix (A25742, Thermo-Fisher) and primers provided in **Table S10**. QuantStudio 5 Flex Real-Time PCR System (Applied Biosystems, Foster City, CA) was used for amplification and fluorescence detection. PCR was performed under the following conditions: 95□ for 10 min, followed by 40 cycles of 95 □ for 15 sec and 60 □ for 1 min. Relative mRNA expression was calculated using the delta-delta Ct method. *Gapdh* was used as a reference transcript.

### RNA-seq analysis

Transcript profiles were generated from wild type and *Akt1*^-/-^, and *Foxo4^Xm-^* junctional zone tissues, and rat differentiated TS cells expressing control or *Foxo4* shRNAs. Complementary DNA libraries from total RNA samples were prepared with Illumina TruSeq RNA preparation kits according to the manufacturer’s instructions (Illumina, San Diego, CA). RNA integrity was assessed using an Agilent 2100 Bioanalyzer (Santa Clara, CA). Barcoded cDNA libraries were multiplexed onto a TruSeq paired-end flow cell and sequenced (100-bp paired-end reads) with a TruSeq 200-cycle SBS kit (Illumina). Samples were run on an Illumina NovaSeq 6000 sequencer at the KUMC Genome Sequencing Facility. Reads from *.fastq files were mapped to the rat reference genome (Ensembl Rnor_5.0.78) using CLC Genomics Workbench 12.0 (Qiagen, Germantown, MD). Transcript abundance was expressed as transcript per million mapped reads (TPM) and a *P* value of 0.05 was used as a cutoff for significant differential expression. Statistical significance was calculated by empirical analysis of digital gene expression followed by Bonferroni’s correction. Pathway analysis was performed using Database for Annotation, Visualization, and Integrated Discovery (**DAVID; Huang *et al*., 2009**).

### Immunohistochemistry

Placentation sites were embedded in optimum cutting temperature (**OCT**) compound and sectioned at 10 μm thickness. Sections were fixed in 4% paraformaldehyde, washed in phosphate buffered saline (pH 7.4) three times for five min each, blocked with 10% normal goat serum (50062Z, Thermo-Fisher), and incubated overnight with primary antibodies: pan cytokeratin (1:300 dilution, F3418, Sigma-Aldrich) to identify trophoblast cells, vimentin (1:300 dilution, sc-6260, Santa Cruz Biotechnology) to distinguish placental compartments, and FOXO4 (1:300 dilution, 21535-1-AP, Proteintech, Rosemont, IL), phospho-Ser^262^ FOXO4 (1:300 dilution, ab126594, Abcam, Cambridge, MA). After washing with phosphate buffered saline (pH 7.4), sections were incubated with corresponding secondary antibodies: Alexa 568-conjugated goat anti-rabbit IgG (1:500 dilution, A11011, Thermo-Fisher) or Alexa 568-conjugated rabbit anti-mouse IgG (1:500 dilution, A9044, Sigma-Aldrich) for 2 h at room temperature. Sections were then stained with DAPI (1/25,000 dilution, D1306, Invitrogen) and mounted using Fluoromount-G mounting media (0100-01, Southern Biotech, Birmingham, AL) and examined microscopically. Fluorescence images were captured on a Nikon 90i upright microscope (Nikon) with a Photometrics CoolSNAP-ES monochrome camera (Roper). The area occupied by cytokeratin-positive cells (invasive trophoblast cells) within the uterine-placental interface was quantified using ImageJ software, as previously described (**Nteeba *et al*., 2020**).

### In situ hybridization

Distributions of transcripts for *Foxo4* and *Prl7b1* were determined on cryosections of rat placentation sites. RNAscope Multiplex Fluorescent Reagent Kit version 2 (Advanced Cell Diagnostics, Newark, CA) was used for in situ hybridization analysis. Probes were prepared to detect *Foxo4* (NM_001106943.1, 1038981-C1, target region: 750-1651) and *Prl7b1* (NM_153738.1, 860181-C2, target region: 28-900). Fluorescence images were captured on a Nikon 80i upright microscope (Nikon) with a Photometrics CoolSNAP-ES monochrome camera (Roper).

### Rat TS cell culture

Blastocyst-derived rat TS cells (**Asanoma *et al*., 2011**) were cultured in Rat TS Cell Medium [RPMI 1640 medium (11875093, Thermo-Fisher), 20% (vol/vol) fetal bovine serum (**FBS**, Thermo-Fisher), 100 μM 2-mercaptoethanol (M7522, Sigma-Aldrich), 1 mM sodium pyruvate (11360-070, Thermo-Fisher), 100 μM penicillin and 50 U/mL streptomycin (15140122, Thermo-Fisher)] supplemented with 70% rat embryonic fibroblast (REF)-conditioned medium prepared as described previously (Asanoma *et al*., 2011), 25 ng/ml fibroblast growth factor 4 (**FGF4**; 100-31, Peprotech), and 1 μg/mL heparin (H3149, Sigma-Aldrich). For induction of differentiation, rat TS cells were cultured for 15 days in rat TS medium containing 1% (vol/vol) FBS without FGF4, heparin, and REF-conditioned medium. In some experiments, rat TS cells were exposed to a phosphatidylinositol 3-kinase (**PI3K**) inhibitor (LY294002, 10 μM, 9901, Cell Signaling Technology), chloroquine (50 μM for 24 h, C6628, Sigma-Aldrich), or staurosporine (1μM for 3 h, 9953, Cell Signaling Technology) followed by Western blot analysis or immunohistochemistry.

### Lentivirus construction and production

Lentivirus construction and production were described previously (**Muto *et al*., 2021; Varberg et al., 2021**). Briefly, the lentivirus encoding the shRNA targeting *Foxo4* was constructed using a pLKO. 1 vector. shRNA oligo sequences used in the analyses are provided in **Table S11**. Lentiviral packaging vectors were obtained from Addgene and included pMDLg/pRRE (plasmid 12251), pRSV-Rev (plasmid 12253), pMD2.G (plasmid 12259). Lentiviral particles were produced using Attractene (301005, Qiagen) in human embryonic kidney (HEK) 293FT (Thermo-Fisher) cells.

### Lentiviral transduction

Rat TS cells were incubated with lentiviral particles for 24 h followed by selection with puromycin dihydrochloride (5 μg/mL; A11138-03, Thermo-Fisher) for two days. Cells were then cultured for 1-3 days in Rat TS Cell Medium prior to differentiation.

### Statistical analysis

Student’s *t*-test, Welch’s *t*-test, Dunnett’s test, or Steel test were performed, where appropriate, to evaluate the significance of the experimental manipulations. Results were deemed statistically significant when *P* < 0.05.

## ACKNOWLEDGEMENTS

The research was supported by postdoctoral fellowships from the Kansas Idea Network of Biomedical Research Excellence, P20 GM103418 (A.M-I.), Lalor Foundation (K.K., A.M-I., M.M.), American Heart Association (K.K., M.M.), and an NIH National Research Service Award, HD104495 (R.L.S.) and NIH grants (HD020676, HD079363, HD099638, HD105734), and the Sosland Foundation. We also thank Stacy Oxley and Brandi Miller for administrative assistance.

## AUTHOR CONTRIBUTIONS

K.K., A.M.-I., and M.J.S. conceived and designed the research; K.K., A.M.-I., K.I., M.-L.W., R.L.S., M.E.S., M.M. performed experiments; K.K., A.M.-I., K.I., M.R.P., and M.J.S. analyzed the data and interpreted results of experiments; K.K., A.M.-I., and M.J.S. prepared figures and manuscript; All authors read, contributed to editing, and approved the final version of manuscript.

## CONFLICT OF INTEREST

There is no conflict of interest that could be perceived as prejudicing the impartiality of the research reported.

## DATA AVAILABILITY

RNA-seq datasets are available at the Gene Expression Omnibus (**GEO**) database, https://www.ncbi.nlm.nih.gov/geo/ (accession number GSE205831). All data generated and analyzed in this study are included in the published article and supporting files. Resources generated from the research are available from the corresponding author upon reasonable request. AKT1 and FOXO4 mutant rat models are available through the Rat Resource and Research Center (Columbia, MO).

## Notes

### Competing Interest Statement

The authors have declared no competing interest.

### Summary of Updates

Results section updated Supplemental files updated

