## Supplementary Data for "AKT1-FOXO4 AXIS RECIPROACLLY REGULATES HEMOCHORIAL PLACENTATION"

**[2]Department of Obstetrics and Gynecology, University of Kansas Medical Center, Kansas City, KS**

**[3]Center for Perinatal Research, Children's Mercy Research Institute, Children's Mercy, Kansas City, MO**

***Contributed equally**

[§]**Present address:** Department of Obstetrics and Gynecology, University of Missouri-Kansas City School of Medicine, Kansas City, MO

[†]**Present address:** Department of Obstetrics and Gynecology, University Hospital, Case Western Reserve University, Beachwood, OH 44122

[‡]**Present address:** Department of Stem Cells and Human Disease Models, Research Center for Animal Life Science, Shiga University of Medical Science, Seta, Tsukinowa-cho, Otsu, Shiga 520-2192, Japan

**This PDF file includes:**
Supplementary Figures 1-4
Supplementary Tables 1, 4, 9-11

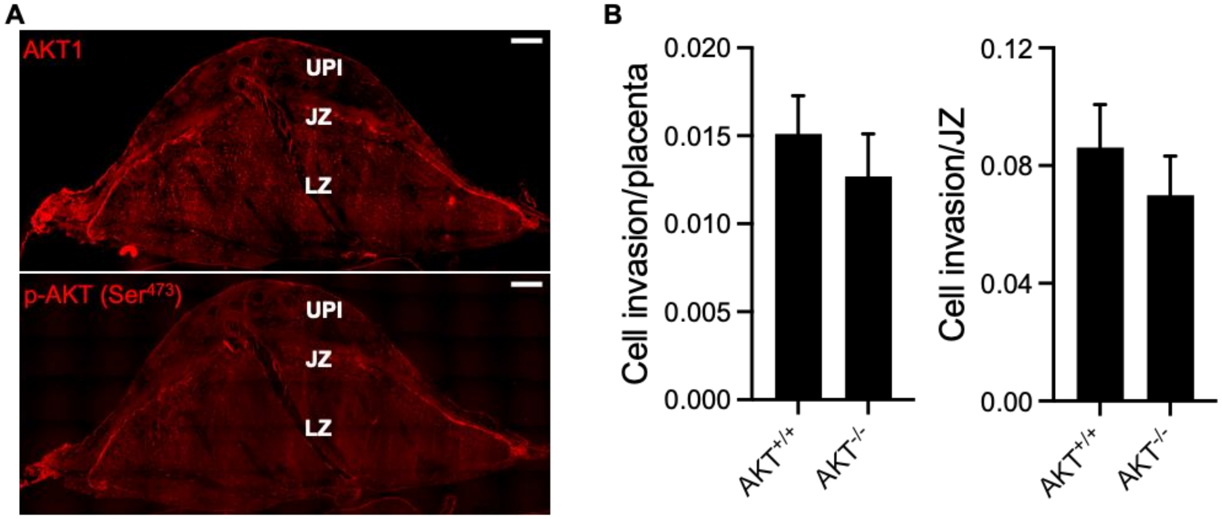


**Figure S1**. **Placental AKT protein distribution and intrauterine trophoblast cell invasion in wild type and *Akt1* null placentation sites**. **A)** Distribution of AKT1 (top) and phospho (p)-AKT4 (Ser$^{473}$) proteins (bottom) in the gestation day (**gd**) 18.5 placentation site. **B**) Area occupied by invasive trophoblast cells (cytokeratin-positive cells) within the uterine-placental interface was quantified and is presented relative to areas associated with the entire placenta or the junctional zone (**JZ**). Graphs represent means ± SEM (n = 6/group).

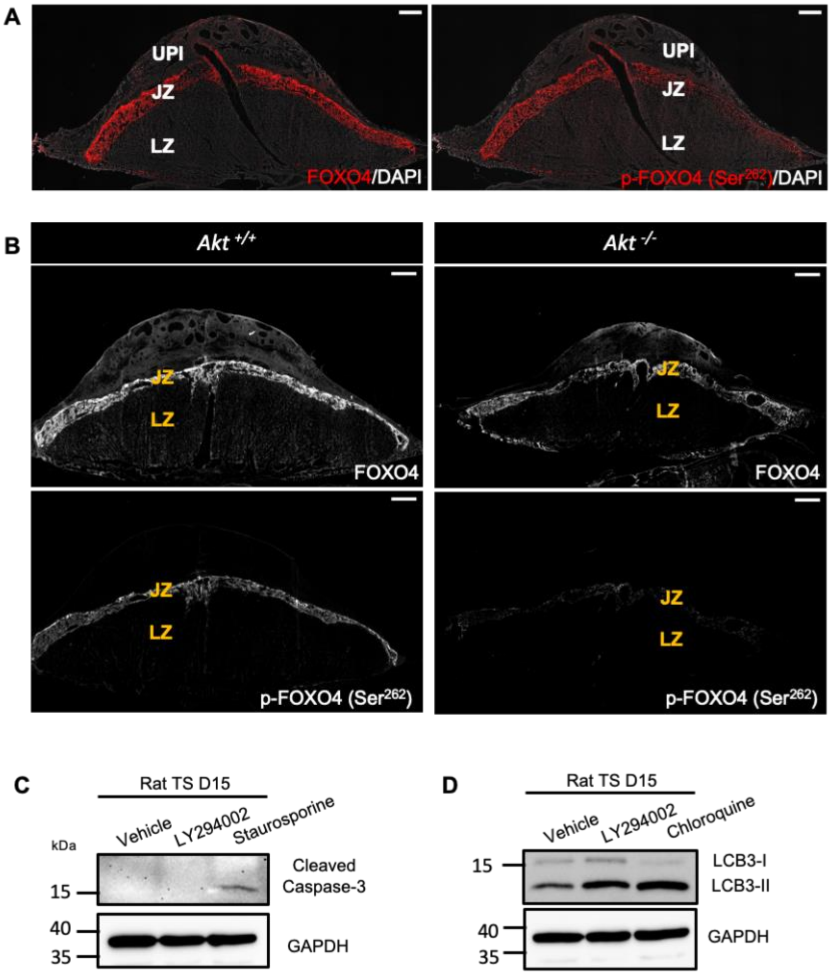


**Figure S2**. **Relationship of PI3K/AKT signaling to FOXO4 distribution and functions. A)** Distributions of FOXO4 and phospho (p)-FOXO4 ($Ser^{262}$) proteins in gestation day (**gd**) placentation sites. **B)** Distribution of FOXO4 and phospho (p)-FOXO4 ($Ser^{262}$) proteins in $Akt1^{+/+}$ and $Akt1^{-/-}$ placentas at gd 18.5. UPI: uterine-placental interface; JZ: junctional zone; and LZ: labyrinth zone. **C**) Western blot analysis of cleaved caspase-3 in rat TS cells exposed to vehicle (DMSO), LY294002 (10 µM), or staurosporine (1 µM). Staurosporine was used as a positive control. **D**) Western blot analysis for serine palmitoyltransferase long chain base subunit 3 (**LCB3**) in rat TS cells exposed to vehicle (DMSO), LY294002 (10 µM), or chloroquine (50 µM). Chloroquine was used as a positive control. Please note that inhibition of PI3K/AKT signaling did not have a detectable effect on the formation of cleaved caspase-3 but did have a modest effect on LCB3 accumulation.

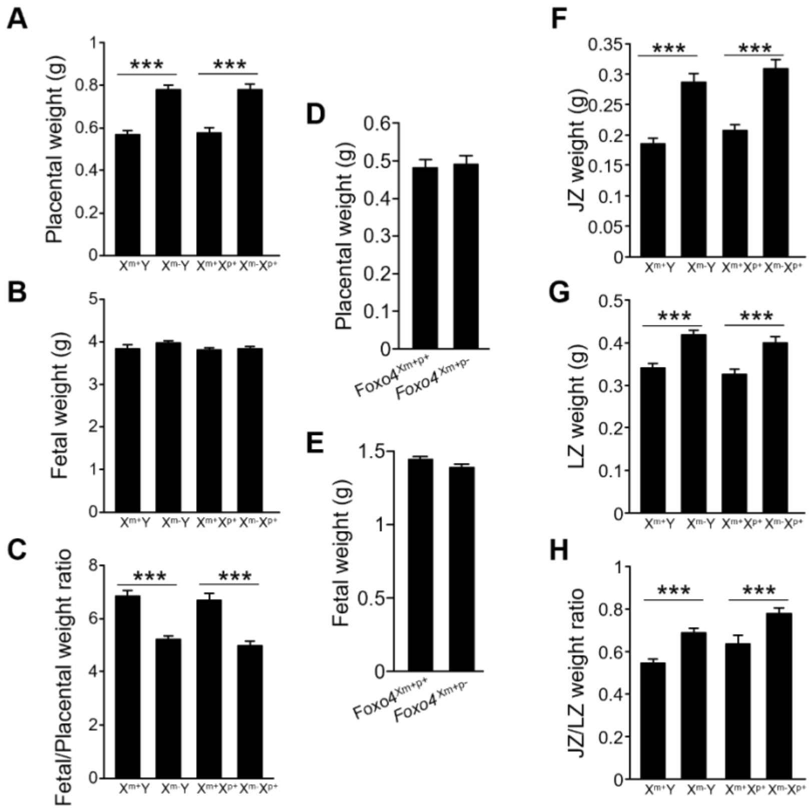


**Figure S3**. Placentas **(A)** and fetuses **(B)** were dissected from *Foxo4* heterozygous females mated with wild type males at gd 20.5 and weighed; **C**, fetus/placenta ratio. Placentas **(D)** and fetuses **(E)** were dissected from wild type females mated with *Foxo4* hemizygous null males at gd 18.5 and weighed. **F-H**) Placentas from heterozygous females mated with wild type males were then separated into junctional zone (**JZ, F**) and labyrinth zone (**LZ, G**) compartments, and weighed; **H**, JZ/LZ weight ratio. $X^{m+}Y$, n = 19; $X^{m-}Y$, n = 22; $X^{m+}X^{p+}$, n = 15; $X^{m-}X^{p+}$, n = 15 from 6 dams. Graphs represent means ± SEM. Asterisks denote statistical differences (***$P$<0.001) as determined by Student's or Welch's *t*-test.

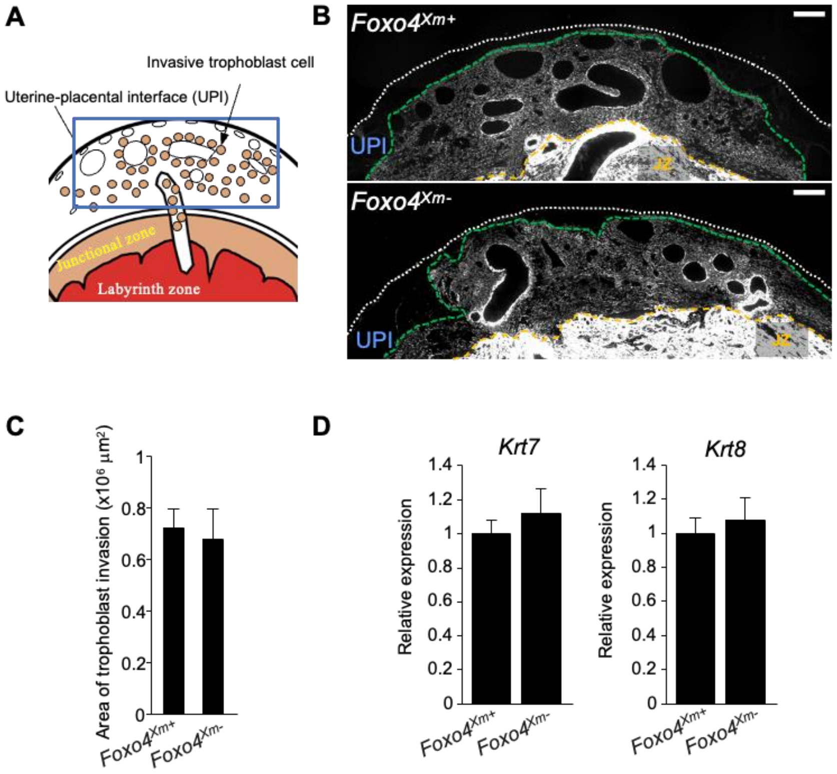


**Figure S4. FOXO4 deficiency does not affect either intrauterine trophoblast invasion or the intrauterine invasive trophoblast cell phenotype. A)** Schematic representation of a late gestation placentation site. The uterine-placental interface, site for intrauterine trophoblast invasion, is highlighted in the boxed area. **B)** Trophoblast cells were immunostained for pan-cytokeratin (**KRT**). Representative images are shown. The extent of intrauterine trophoblast invasion is demarcated using a green dashed line. The white dotted line represents the outer border of the uterus, and the yellow dashed line represents the uterine border with the placenta. Scale bars = 500 µm. **C)** The area of intrauterine trophoblast invasion is graphically depicted (n = 6/group). Graphs represent means ± SEM. **D)** RT-qPCR measurements of *Krt7* and *Krt8* transcripts, signature markers for invasive trophoblast cells, within dissected uterine-placental interface tissue specimens at gd 18.5 (*Foxo4*$^{Xm+}$, n = 12; *Foxo4*$^{Xm-}$, n = 12). Graphs represent means ± SEM.

**Table S1. Genotype of offspring following *Akt1* heterozygous breeding.**

| **Offspring from *Akt1*$^{+/-}$ x *Akt1*$^{+/-}$** | **Offspring genotype** | | |
|---|---|---|---|
| | ***Akt1*$^{+/+}$** | ***Akt1*$^{+/-}$** | ***Akt1*$^{-/-}$** |
| Total | 40 | 92 | 42 |
| Ratio (%) | 23 | 52.9 | 24.1 |

**Table S4. Genotype of offspring following *Foxo4* hemizygous male x wild type female breeding.**

| **Offspring from $X^{-}Y$ x XX** | **Offspring genotype** | |
|---|---|---|
| | **$X^{m+}Y$** | **$X^{m+}X^{p-}$** |
| Total | 40 | 44 |
| Ratio (%) | 47.6 | 52.4 |

**Table S9. Primers used for genotyping**

| Primer name | Sequence |
|---|---|
| *Akt1* Fwd | TGAGTCCATTCTGGAGGACTAGAC |
| *Akt1* Rev1 | TTGCCCAGTAGCTTCAGGTACTC |
| *Akt1* Rev2 | GAGGGAAGGTTAGGGACTAGCC |
| *Foxo4* Fwd | AGAAGGTACCCACGGAGGGA |
| *Foxo4* Rev1 | CCACACAGTTCCTGCTGTACATAG |
| *Foxo4* Rev2 | CTCCTTGGAGTGGCACCTTC |

Fwd, forward
Rev, reverse

**Table S10. List of primers used for RT-qPCR.**

| Gene | Forward primer | Reverse primer | Accession no. | Amplicon size (bp) |
|---|---|---|---|---|
| ***Ccn3*** | CATGGTTCGGCCTTGTGAG | TGGATTTCAGGGACTTCTTGGT | NM_030868.2 | 89 |
| ***Ccnd1*** | AATGCCAGAGGCGGATGAGA | CGTTGTGCGGTAGCAGGAGA | NM_171992.5 | 190 |
| ***Ccne1*** | TGCAGGCGAGGATGAGA | GAAGTCCTGTGCCAAGTAGAATG | NM_001100821.1 | 98 |
| ***Cdc6*** | CAGGCGAGCTATTGAAATTGTG | GACTTGGGATATGTGAGCGAGA | NM_001108298.1 | 130 |
| ***Cdk1*** | GTTGACATCTGGAGCATAGG | CTCTACTTCTGGCCACACTT | NM_019296.2 | 144 |
| ***Mcm5*** | TGTCCAGGATTTCACCAAACA | CACTTGAGGCGGTAAAGCAC | NM_001399204.1 | 123 |
| ***Prl8a4*** | CTGAAACCCTCTGTAATCTTGCTG | GTCTCGTCCCTCTTAATCAGTTTTG | NM_021580.1 | 112 |
| ***Krt7*** | CGGAATGGAACCTGTGAA | GTAGATGTAGTCTTGATGGAATAAG | NM_001047870.2 | 150 |
| ***Krt8*** | TGGGCCAGGAGAAGCTGAA | CACATCCTTGATGAGGACAAA | NM_199370.1 | 140 |
| ***Foxo4*** | CGGAATGCCTGGGGAAA | ATGTACCTTGATGAACTTGCTGTG | NM_001106943.1 | 213 |
| ***Grb7*** | TACCACCTGGAGAGAAGAGAGAGAG | GGGCTCAGATCCAGTTCCA | NM_053403.2 | 141 |
| ***Erbb3*** | TGCGTTGCCAGTTGTCC | CCGTGCTTATCTACTTCCATCTTGT | NM_017218.3 | 92 |
| ***JamL*** | TCGGCCTTGATGGGATG | CACGCTGAGGCTGGAGTAGTAG | XM_032909807.1 | 129 |
| ***Lpal2*** | AAGGAGATGCCAACCAACAAA | GCCATTCTTCCCTCTCCTGA | NM_001109578.1 | 125 |
| ***Mmp12*** | GCTGGTTCGGTTGTTAGG | GTAGTTACACCCTGAGCATAC | NM_053963.2 | 100 |
| ***Gstm1*** | CTGGACGCCTTCCCAAA | TAGCAAGGGCCTACTTGTTACTCC | NM_017014.2 | 145 |
| ***Txnip*** | GTCTCAGCAGTGCAAACAGACC | AAGCTCAAAGCCGAACTTGTACTC | NM_001008767.2 | 139 |
| ***Alox5ap*** | TGTGGGCAATGTTGTGCTC | GCTTTGCGCCTTGCTTTC | NM_017260.2 | 100 |
| ***Nupr1*** | GCCCACACTTCCCAGCA | ACCTCCACCGACGACATAAGA | NM_053611.2 | 102 |
| ***Gapdh*** | GACATGCCGCCTGGAGAAAC | AGCCCAGGATGCCCTTTAGT | NM_017008.4 | 92 |

**Table S11. shRNA sequences**

| shRNA | Sequence |
| --- | --- |
| *Foxo4* shRNA 1 | CCGGTGCAGTCCTTGTCCTCGAAACTCGAGTTTCGAGGACAAGGACTGCTTTTTG |
| *Foxo4* shRNA 2 | CCGGTGCTTGCATCTCCTACTGAACTCGAGTTCAGTAGGAGATGCAAGCTTTTTG |
